# Luminal epithelium remodeling underlies endometrial regeneration during menstruation and pregnancy

**DOI:** 10.64898/2026.03.08.710375

**Authors:** Claire J. Ang, Jaina L. R. Gable, Kayla C. Lyons, Enya Miguel Whelan, Çağrı Çevrim, Taylor D. Skokan, Patricia L. Murphy, Sofia G. Bennetts, Benjamin D. Manetta, Aellah M. Kaage, Dhanvika Mopure, Ana Breznik, Allison E. Goldstein, Fátima Sanchís-Calleja, Thomas E. Spencer, Andrew M. Kelleher, Kara L. McKinley

## Abstract

Menstruation and pregnancy impose an immense regenerative burden on the endometrium. These events breach the luminal epithelium lining the uterine cavity, which is proposed to be replenished by cells in adjoining epithelial glands. In contrast to this gland-centric model, we find that luminal and glandular epithelia are maintained by separate progenitor populations during homeostasis, induced menstruation, pregnancy, and postpartum repair in mice. Although our data indicate that gland cells do not contribute substantially to luminal epithelium regeneration under physiological conditions, we find that they can serve as facultative progenitors that resurface the tissue after chemical ablation. During menstruation, the luminal epithelium bypasses the need for gland contributions by undergoing extensive expansion and morphogenesis to re-epithelialize stromal surfaces concurrently with tissue breakdown. Analogous morphogenesis occurs during gestation, revealing luminal epithelial expansion as a unifying mechanism enabling simultaneous stromal disruption and re-epithelialization, which may underlie the endometrium’s remarkable regenerative capacity.

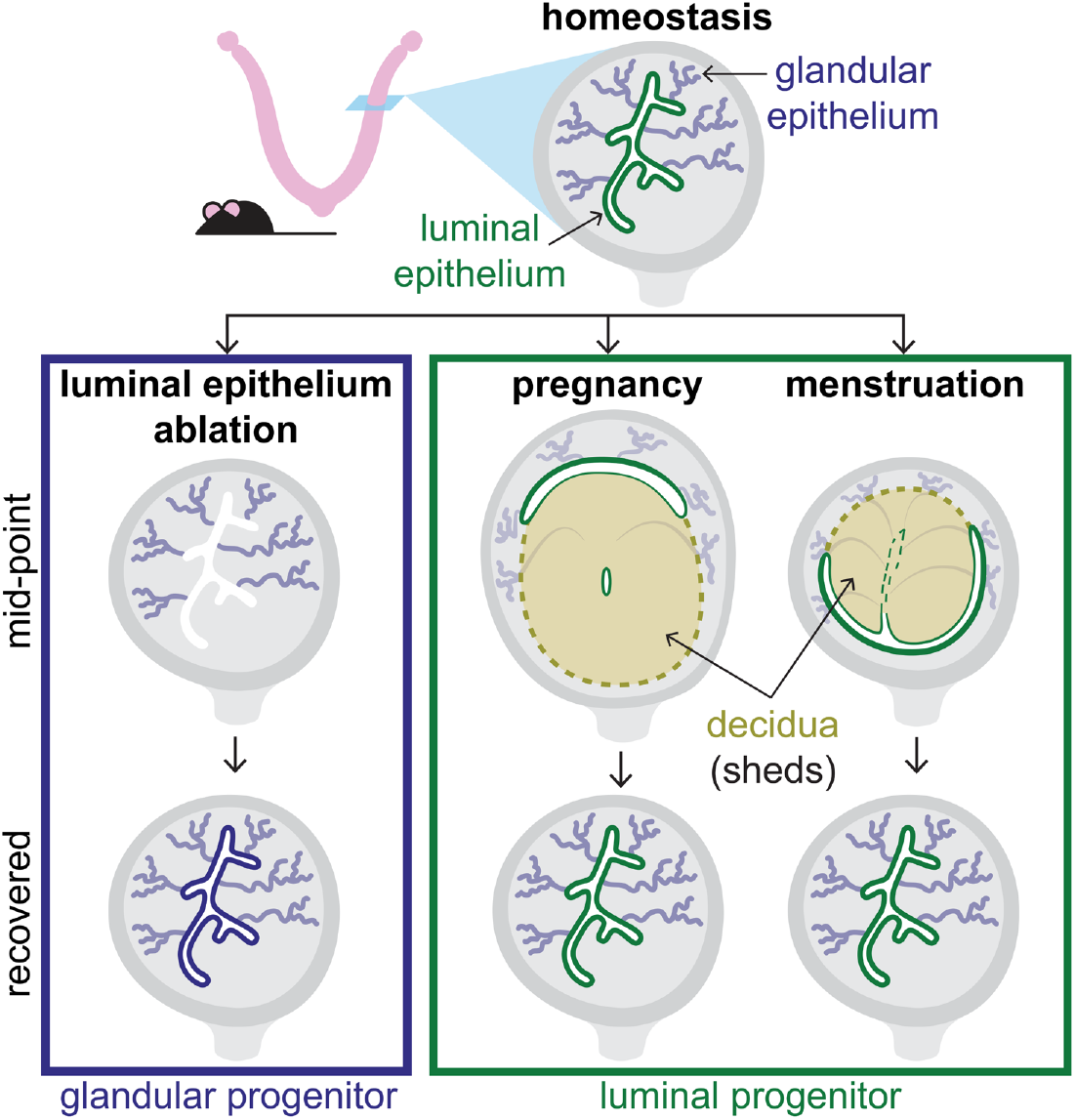

## Introduction

The endometrial tissue that lines the uterine cavity undergoes dramatic remodeling and disruption during menstruation, pregnancy and childbirth (parturition). During the human menstrual cycle, hormone changes stimulate the endometrium to differentiate into a specialized tissue called the decidua. If pregnancy occurs, the decidua persists and remodels around the embedding embryo and its developing placenta until delivery. In the absence of pregnancy, the decidua is shed during menstruation. Each of these events perturbs the endometrium’s major structural components, including a luminal epithelium, branched epithelial glands, and the surrounding stroma (Figure 1A). Proper restoration of the epithelium is crucial for fertility, as the luminal epithelium serves as a mediator of decidualization and the site of initial embryo attachment,^1,2^ and the glands secrete factors to support early pregnancy.^3-5^ However, our understanding of endometrial epithelial regeneration remains incomplete.

**Figure 1.**
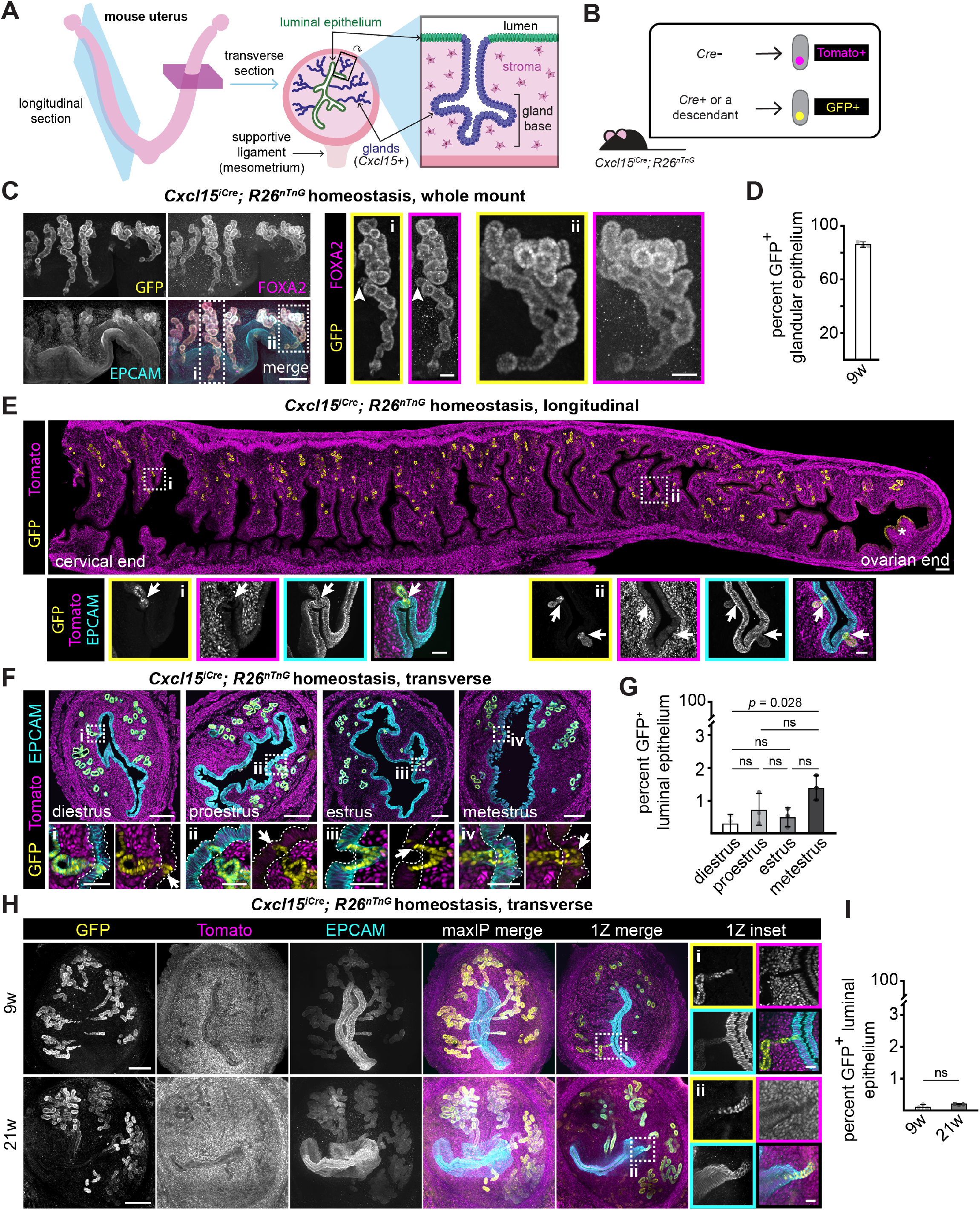
The *Cxcl15* gland lineage makes minimal contributions to the endometrial luminal epithelium in homeostasis. (A) Schematic of mouse uterus indicating transverse (200 µm thick) and longitudinal (50 µm thick) section orientations. All samples in the manuscript are presented with the mesometrium down. (B) Schematic of fluorescent reporter expression in *Cxcl15*^*iCre*^; *R26*^*nTnG*^ mice. (C) Representative immunofluorescence image of cleared whole mount uteri collected during estrus. Arrowhead indicates a region with GFP-negative/FOXA2-positive nuclei. (D) Quantification of the proportion of FOXA2-positive (gland) nuclei expressing GFP in homeostatic uteri of 9 w old *Cxcl15*^*iCre*^; *R26*^*nTnG*^ mice. n = 3. (E) Representative immunofluorescence image of the entire length of a uterine horn collected during estrus. Asterisk indicates oviduct epithelium. (F) Representative immunofluorescence images of 12 µm-thick cross-sections of uteri collected during each estrous cycle stage. Dotted lines indicate luminal epithelium as defined for quantification. (G) Quantification of the proportion of luminal epithelium expressing GFP in *Cxcl15*^*iCre*^; *R26*^*nTnG*^ uteri collected at each estrous cycle stage. (H) Representative immunofluorescence images of uteri collected in homeostasis, at 9 or 21 w of age. Brightness/contrast were adjusted independently for each sample. (I) Quantification of the proportion of luminal epithelium nuclei positive for GFP in uteri collected in homeostasis, at 9 or 21 w of age. For all images, single channel outsets are shown are as maxIPs. Inset channel is indicated by box color. Arrows indicate GFP-positive junctions between glands and luminal epithelium. For all graphs, each dot represents an individual animal and error bars represent standard deviation. Outset scale bars, 200 µm; inset scale bars, 50 µm.

The bases of the glands have been proposed as a key reservoir for endometrial epithelial progenitor cells in humans. This model is particularly appealing in the context of menstruation. Electron microscopy studies of the menstruating human uterus have shown that shedding of the superficial layer of the endometrium (the functionalis) exposes an underlying endometrial layer (the basalis), composed of gland fragments and remnant stroma.^6-9^ The basalis glands are enriched in putative stem cell markers and features associated with stemness, such as the capacity for clonogenicity and self-renewal *in vitro*.^10-15^ One study examining the human endometrium found that spontaneously arising mutant clones in the basalis glands expand to the functionalis glands with age, supporting a role for basalis progenitors in regenerating the functionalis glands.^16^ Whether these cells can generate luminal epithelium has not been extensively assessed, although recent work has revealed that glandular cells can adopt aspects of luminal epithelial gene expression when disrupted.^17,18^

Alongside the ongoing work to identify human endometrial epithelial progenitors, several studies have performed lineage tracing in mice to dissect the roles of glandular epithelial cells during endometrial homeostasis and regeneration.^19-21^ The findings of these studies have varied. Lineage tracing of gland populations using genetic drivers ubiquitously expressed in glands (*Lgr5* and *Foxa2*) revealed no contribution to the luminal epithelium during adulthood, even after 6 weeks or 26 weeks, respectively.^20,21^ In contrast, a subset of glandular cells expressing *Axin2* has been found to contribute to the adult luminal epithelium over a 12-week span.^19^ The effect of menstruation on progenitor dynamics was not assessed in these studies. Mice do not menstruate naturally, although menstruation-like endometrial shedding can be induced through a variety of methods combining experimental hormone control with the addition of a stimulus to induce the formation of an artificial decidua (sometimes called a deciduoma), which sheds upon hormone withdrawal.^22^ Mouse models of menstruation and other injuries provide an opportunity to apply genetic lineage tracing to assess whether glandular epithelium contributions vary across physiological contexts.

Using a *Cxcl15-Cre* mouse model to specifically track gland cells and their descendants, we found that *Cxcl15-*lineage gland cells make minimal contributions to the luminal epithelium during homeostasis, induced menstruation, pregnancy, and parturition, but are competent to re-epithelialize the lumen following extensive loss of luminal epithelium through chemical ablation. During induced menstruation, we observed persistent sheets of luminal epithelium that envelop the decidua and re-epithelialize the lumen concurrently with tissue shedding, bypassing the need for gland contributions. We identified analogous luminal epithelium morphogenesis occurring around the decidua and fetal tissues during gestation. These findings reveal luminal epithelial expansion concomitant with tissue disruption as a unifying mechanism by which the uterus achieves rapid epithelial restoration in menstruation and pregnancy.

## Results

### The *Cxcl15*-labeled gland lineage contributes minimally to the luminal epithelium during homeostasis

We first sought to test the contributions of gland progenitors to unperturbed luminal epithelium using the *Cxcl15*^*iCre*^; *R26*^*nTnG*^ mouse line, which induces a permanent switch from TdTomato to GFP expression in *Cxcl15-*expressing cells and their descendants (Figure 1B).^23^ *Cxcl15* expression is specifically detected in the glandular epithelium starting on postnatal day 12, when the glands first develop from the luminal epithelium. In contrast, expression is not detected in the luminal epithelium.^23^ In young adults (9 weeks old), 86% of cells expressing the gland marker, *Foxa2*, were GFP-positive, indicating robust labeling of glandular epithelial cells (Figure 1C and 1D). In contrast to the glands, no more than 1.38% of the luminal epithelium was GFP-positive at any estrous cycle stage; instead, the luminal epithelium was predominantly tdTomato-positive/GFP-negative (Figure 1E-G and Table S1). Most of the GFP-positive cells resided at the junctions between glands and luminal epithelium (Figure 1E and 1F insets). Although luminal epithelial cells proliferate every 3 days,^19^ the rare GFP-positive cells did not expand long term: only 0.09% and 0.19% of luminal epithelial cells expressed GFP at 9 and 21 weeks of age, respectively (Figure 1H, 1I and Table S1). These data demonstrate that the *Cxcl15* gland lineage makes minimal contributions to the luminal epithelium during homeostasis.

### The *Cxcl15*-labeled gland lineage regenerates the luminal epithelium after extensive epithelial ablation

Although *Cxcl15*-lineage labeling remained restricted to the glands during homeostasis, we considered whether tissue disruption might enable cross-compartment plasticity, as occurs in other epithelial systems, including the skin.^24,25^ To test this, we induced extensive loss of luminal epithelium using transient exposure to the liquid surfactant, polidocanol, which is used as a model of epithelial damage in the airway (Figure 2A).^26^ By 2 h post-ablation, nearly all of the luminal epithelium and gland junctions directly abutting the uterine cavity were ablated, while underlying glandular epithelium remained intact (Figure 2B). By 72 h post-ablation, the luminal epithelium had regenerated (Figure 2B). To assess whether the repaired luminal epithelium was gland-derived, we performed ablation experiments in *Cxcl15*^*iCre*^; *R26*^*nTnG*^ mice. 72 h after ablation, the luminal epithelium was almost entirely GFP-positive throughout the uterus (5/6 animals; Figure 2C and 2D). Most luminal epithelial cells remained GFP-positive 14 and 28 days after polidocanol treatment, suggesting long-term persistence (Figure S1A and S1B). Together, our data indicate that *Cxcl15*-lineage cells can resurface the endometrium after extensive luminal epithelium ablation.

**Figure 2.**
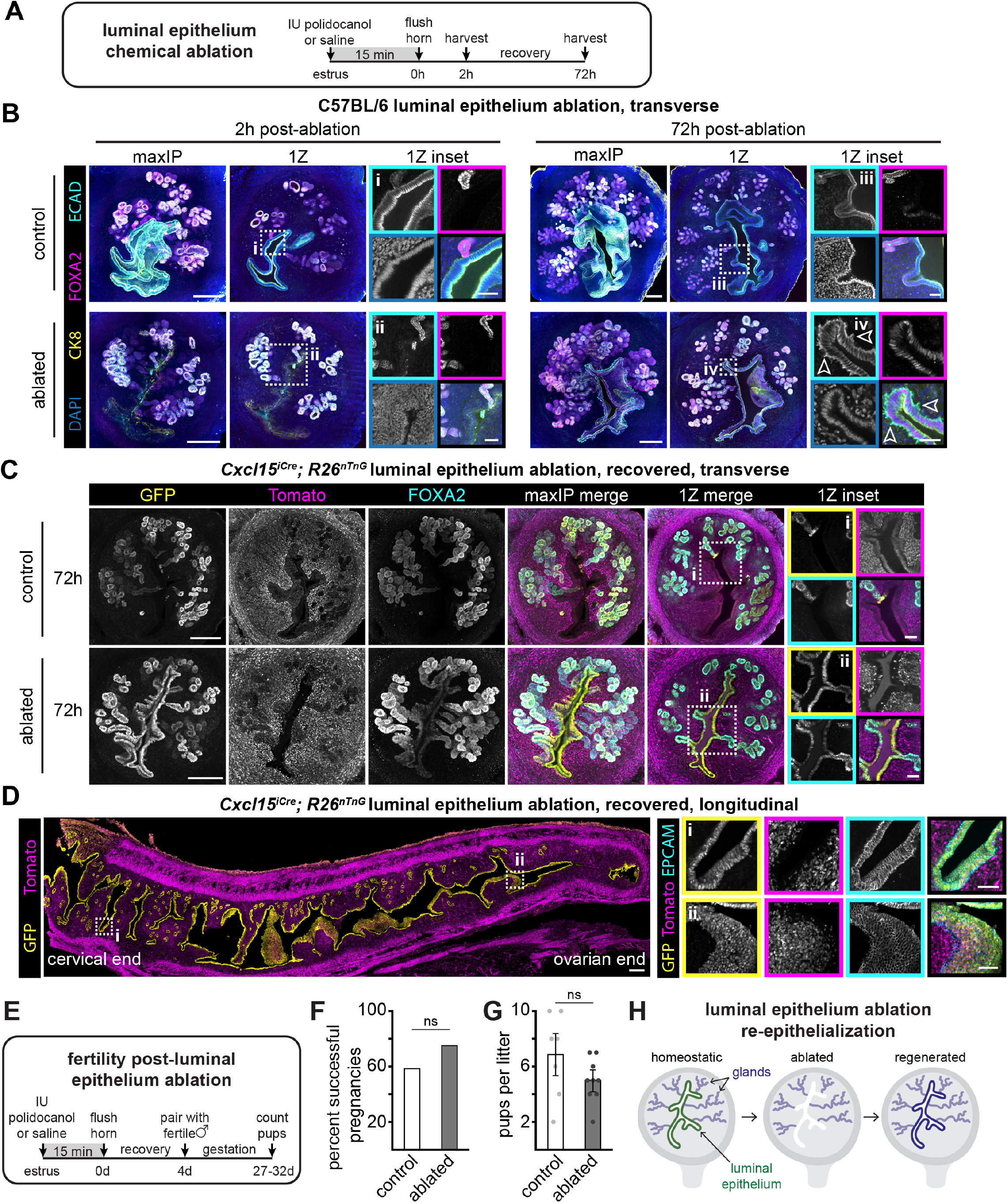
Luminal epithelium ablation stimulates *Cxcl15* gland lineage contributions. (A) Experimental timeline for luminal epithelium chemical ablation. (B) Representative immunofluorescence images of uteri collected as indicated in A. Outlined arrowheads indicate cell membrane protrusions from the basal side of the luminal epithelium. (C) Representative immunofluorescence images of uteri collected as indicated in A. Brightness/contrast were independently adjusted for each single Z plane and inset panel. (D) Representative immunofluorescence image of the entire length of a uterine horn collected 72 h after luminal epithelium ablation. (E) Experimental timeline for testing fertility in C57BL/6 mice following luminal epithelium ablation or saline injection. (F) Quantification of the proportion of C57BL/6 females that carried a litter to term following recovery from ablation or saline injection as indicated in E. n = 12 per condition. (G) Quantification of the number of pups per litter from C57BL/6 dams following recovery from ablation or saline injection as indicated in E. Each dot represents a litter, and error bars represent standard deviation. n = 7 control; n = 9 ablated. (H) Summary schematic of recovery from luminal epithelium ablation. For all images, single channel outsets are shown as maxIPs. Inset channel is indicated by box color. Outset scale bars, 200 µm; inset scale bars, 50 µm. ECAD, E-cadherin (pan-epithelial marker); CK8, cytokeratin 8 (pan-epithelial marker); IU, intrauterine.

Upon initial recovery from polidocanol treatment, the repaired luminal epithelium exhibited transient, basal cell membrane projections (Figure 2B iv). Additionally, we observed persistent, aberrant FOXA2 expression throughout the luminal epithelium, although at lower levels than the glands (Figure 2C ii, S1A and S1B). FOXA2 is canonically restricted to the glands during adulthood and serves critical functions during gland development and fertility.^27,28^ In light of these aberrations, we evaluated the function of luminal epithelium derived from the *Cxcl15* lineage by testing its competence to facilitate pregnancy (Figure 2E). There was no significant difference in the proportion of dams that were able to carry litters to term, or litter size between polidocanol-treated dams and unablated controls (Figure 2F and 2G). Thus, although the *Cxcl15* gland lineage makes minimal contributions to the luminal epithelium during homeostasis, it has the capacity to form functional, albeit altered, luminal epithelium upon extensive chemical ablation (Figure 2H).

### Persistent luminal epithelium covers shedding and retained tissue during induced menstruation

Our data indicated that two different progenitors can replenish the luminal epithelium in a context-dependent manner. In homeostasis, luminal populations appeared to be self-renewing, but after a chemical ablation injury, glandular progenitors contributed to luminal epithelium restoration. We sought to determine which of these progenitor populations contribute in response to physiologically relevant disruptions to the endometrium, beginning with menstruation.

Menstruation is proposed to proceed through shedding of luminal epithelial cells and superficial stromal cells, leaving behind an exposed stromal surface dotted with remnant basalis glands, akin to our chemical ablation model. Thus, we hypothesized that gland progenitors would regenerate the luminal epithelium after induced menstruation. We induced menstrual-like shedding in mice by hormone administration and intrauterine oil injection to trigger decidualization (Figure S2A).^22^ The cessation of hormone administration triggers decidual shedding, which is detectable as blood in vaginal lavages an average of 2 to 4.5 days later. Endometrial re-epithelialization proceeds efficiently despite the absence of ovarian hormones.^29^ We detected substantial apoptotic debris in the stroma and ongoing remodeling shortly after bleeding cessation (Figure S2B). By 10 days after the last progesterone injection, when the tissue had restored its homeostatic architecture, we were surprised to find that only 0.27% of luminal epithelial cells were GFP-positive (Figure 3A, 3B and Table S1). These results indicate that the *Cxcl15* gland lineage does not make a substantial long-lasting contribution to the luminal epithelium after menstrual-like shedding.

**Figure 3.**
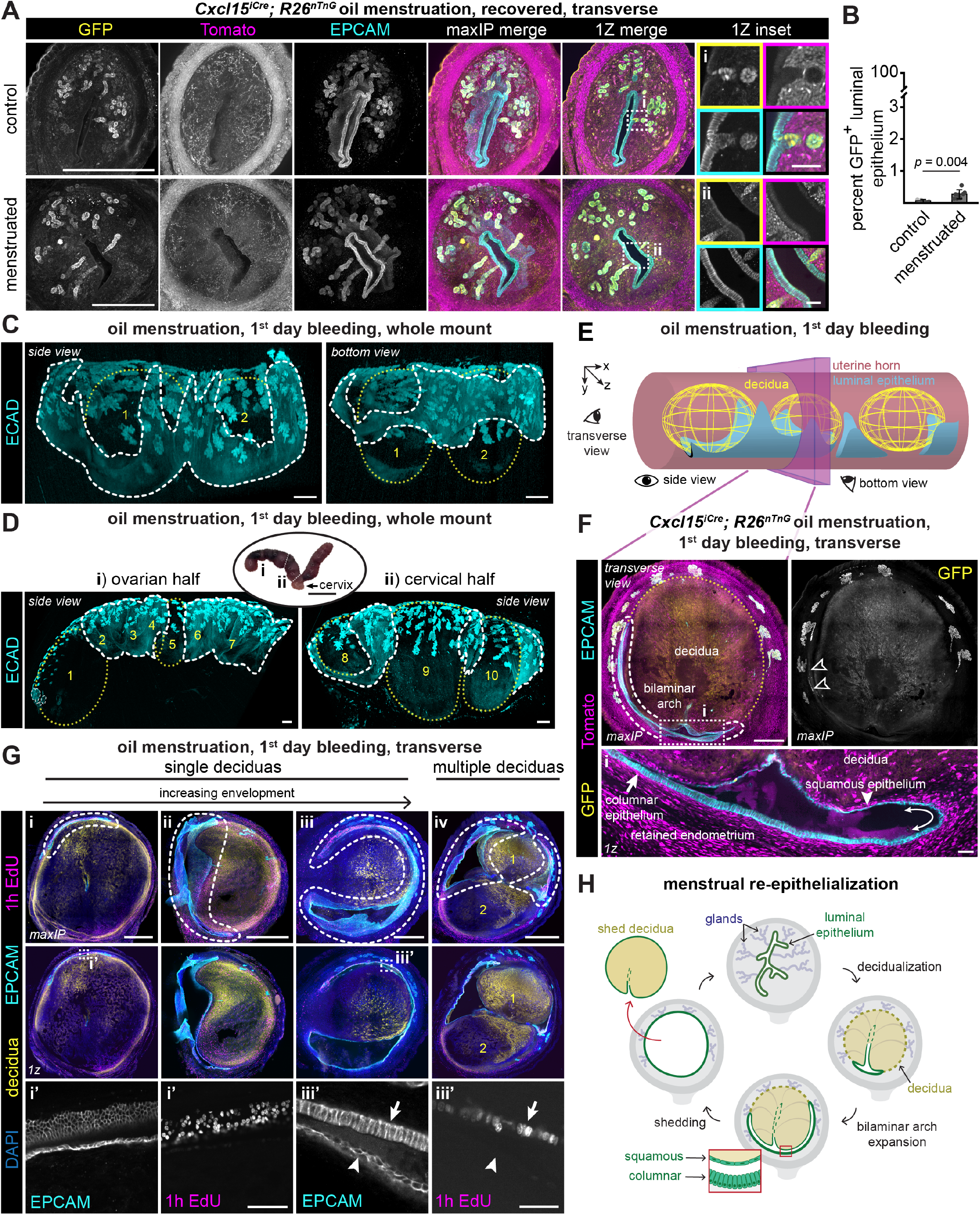
Epithelial sheets envelop decidualized tissue during induced menstruation. (A) Representative immunofluorescence images of uteri collected after recovery from oil-induced menstruation as indicated in Figure S2A paradigm 3. Brightness/contrast were adjusted independently in each sample. (B) Quantification of the proportion of luminal epithelium nuclei positive for GFP in *Cxcl15*^*iCre*^; *R26*^*nTnG*^ uteri after recovery from oil menstruation or saline injection. Each dot represents one animal, and error bars represent standard deviation. n = 4 control; n = 8 menstruated. (C) Representative immunofluorescence images of a cleared whole mount uterine horn (cervical half) collected on the first day of oil menstruation bleeding as indicated in Figure S2A paradigm 1. Side and bottom view perspectives relative to the epithelial sheets are indicated in E. Images were scaled with gamma adjustment and adjusted to maximize opacity. (D) Representative immunofluorescence images of both halves of a cleared whole mount uterine horn collected on the first day of oil menstruation bleeding as indicated in Figure S2A paradigm 1. Gross morphology of whole uterus, and horn division points are shown in ellipse. Images are scaled with gamma adjustment and adjusted to maximize opacity. Ellipse scale bar, 1 cm. (E) Schematic of hypothesized 3-dimensional epithelial sheet structure in relation to decidua. Glands are not shown. Sheets can occur on either the mesometrial or anti-mesometrial side of the horn (compare 3C and 3D). (F) Representative immunofluorescence images of uteri collected on the first day of oil menstruation bleeding as indicated in Figure S2A paradigm 1. Outlined arrowheads indicate GFP-positive luminal epithelial cells. Double ended curved arrows indicate transition between squamous and columnar epithelium. (G) Immunofluorescence images demonstrating variation between uteri collected on the 1^st^ day of oil menstruation bleeding as indicated in Figure S2A paradigm 1. P-cadherin (decidua marker) is shown in yellow. Brightness/contrast were adjusted independently for each sample. (H) Summary schematic of hypothesized model of menstrual re-epithelialization. First, decidualization displaces glands and luminal epithelium to the perimeter of the decidua. Next, proliferation of the columnar epithelium drives bilaminar arch expansion along the detachment plane, covering both the decidua and the basal endometrium. As a result, the shed and retained tissues are almost completely epithelialized by the time of tissue expulsion. For all images, single channel outsets are shown as maxIPs, and insets are shown as single Z planes. Inset channel is indicated by box color unless directly noted. White dashed lines indicate epithelial sheets/bilaminar arch structures, yellow dotted lines and numbering indicate decidual masses, solid arrows indicate basal-side columnar epithelium, solid arrowheads indicate squamous decidual-side epithelium. Outset scale bars, 500 µm, except for panel D ellipse; inset scale bars, 50 µm. ECAD, E-cadherin (pan-epithelial marker).

Based on our lineage tracing results, we considered the possibility that a population of *Cxcl15-*negative luminal epithelial cells may persist during menstruation, providing an alternative cellular reservoir for re-epithelialization. Consistent with this, previous histological studies of induced menstruation have described persistent epithelia with morphologies typical of the luminal epithelium, although their lineage identity and spatial distribution were not assessed.^30^ To extend these observations, we performed tissue clearing and lightsheet imaging to characterize the extent of retained epithelium across the horn on the first day of bleeding.

This menstruation model induces spatially discontinuous decidualization, producing multiple discrete decidual masses along the uterine horn.^31,32^ We observed that each decidual mass pushed the gland bases to the periphery of the tissue, as previously reported.^31,33^ In addition to the glands, we detected broad epithelial sheets overlaying the decidual masses (Figure 3C). The extent of epithelial coverage around decidual masses varied across the uterine horn (Figure 3D). In some regions, continuous sheets overlaid multiple masses (Figure 3D i and Video S1), whereas in other regions the epithelial sheets were interrupted (Figure 3D ii, 3E and Video S2). To interrogate the cell lineages present in these epithelial sheets, we examined thick transverse sections of *Cxcl15*^*iCre*^; *R26*^*nTnG*^ uteri at the same timepoint (Figure 3F and S2C). The epithelial sheets were predominantly GFP-negative, containing only a few rare patches of GFP-positive cells. These data are consistent with a model in which persistent luminal epithelium lines the surface of decidual masses during induced menstruation (Figure 3E).

Intriguingly, the GFP-negative epithelial sheets covered both the retained endometrium and shedding tissue, forming a bilaminar arch that curved around the periphery of the decidua, reminiscent of a smile (Figure 3F). Closer examination revealed that the layers exhibited distinct epithelial morphologies: extending the smile analogy, the decidua-adjacent “upper lip” epithelium was squamous, while the “lower lip” epithelium lining the retained endometrium was columnar (Figure 3F). The transitions between these two layers formed hairpin turns (the “corners of the mouth”) directed towards the sites of delamination between the shedding and retained tissues. Co-staining with the decidual marker, P-cadherin,^34^ revealed that the bilaminar arches encircled decidual masses to different degrees within and between uteri, possibly representing different stages along the progression of decidua envelopment (Figure 3G i, 3G ii and 3G iii). Indeed, a 1 h EdU pulse revealed extensive epithelial proliferation in the columnar layer (Figure 3G i’ and 3G iii’). Together, our data support a model in which proliferation in the basal, columnar layer expands the bilaminar arch along the detachment plane, enabling the epithelium to envelop decidual tissue as shedding proceeds. (Figure 3H).

We detected bilaminar arches in 70% of serial transverse sections collected along the entire horn (76/108 sections, n = 3 mice), consistent with the discontinuous epithelial sheets we observed in our whole mount samples. When adjacent decidual masses were captured in a single section, each was partially encircled with bilaminar arches (Figure 3G iv). We also detected bilaminar arches, albeit more rarely, in an alternative model of menstruation (X-Mens),^34^ which induces decidualization chemogenetically (Figure S2D and S2E). This difference may arise from more extensive or uniform decidualization in the X-Mens model. Thus, the perimenstrual endometrial epithelium adopts a bilaminar arch morphology, with arch formation and the extent of encapsulation varying across the uterine horn and between menstruation models.

### Luminal epithelium envelops decidualized tissue during gestational remodeling and resurfaces the endometrium post-partum

Like menstruation, pregnancy and parturition dramatically alter the endometrial epithelium^31,33,35,36^ but progenitor dynamics during these events remain to be elucidated. To test for gland lineage contributions to the luminal epithelium during early gestation, we evaluated *Cxcl15*^*iCre*^; *R26*^*nTnG*^ dam uteri collected on gestational day 7.5. As in our induced menstruation samples, our analyses revealed extensive GFP-negative, proliferative bilaminar arches surrounding each discrete implantation site, with squamous epithelium lining the decidua, and columnar epithelium lining the basal endometrium (Figure 4A and 4B). We confirmed that these epithelial structures were of maternal rather than fetal origin by lineage tracing fetal tissues (Figure S3A and S3B). Continued expansion of these arches throughout pregnancy^36^ may account for the observation that the tissue is almost completely resurfaced by GFP-negative luminal epithelial cells by the day of parturition, leaving behind only a small placental detachment site (Figure 4C). Indeed, a fortuitous natural case of fetal resorption collected at term revealed that the resorption site was almost completely enveloped in GFP-negative luminal epithelium (Figure S3C). We also observed minimal contribution from GFP-positive cells to placental detachment site re-epithelialization, which was complete within 3 days of parturition (Figure 4C). Even after 4 consecutive pregnancies, GFP-positive cells remained rare in the luminal epithelium of *Cxcl15*^*iCre*^; *R26*^*nTnG*^ mice (Figure 4D). Taken together, these data indicate that luminal epithelium remodeling during and after pregnancy involves minimal contribution from the *Cxcl15* lineage and instead proceeds through the expansion of luminal epithelium with extensive structural similarity to bilaminar arch expansion in menstrual-like shedding (Figure 4E).

**Figure 4.**
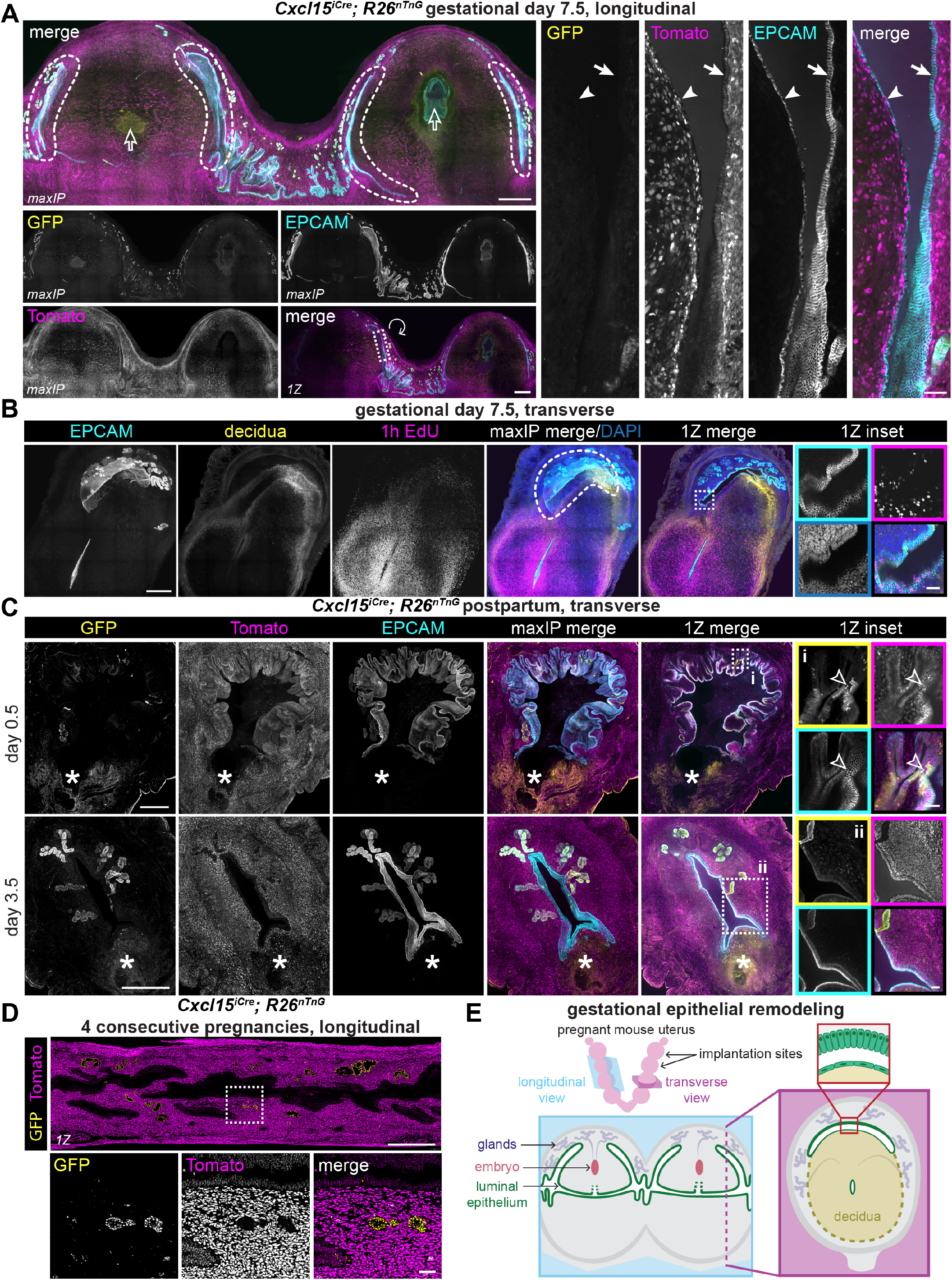
Epithelial sheets envelop decidualized tissue during gestation and restore epithelial integrity during postpartum repair (previous page) (A) Representative immunofluorescence images of two adjacent implantation sites. See E for a representation of section orientations. Outlined arrows indicate embryos, solid arrows indicate basal columnar epithelium, and solid arrowheads indicate squamous decidual-side epithelium. (B) Representative immunofluorescence images of an implantation site collected on gestational day 7.5. P-cadherin (decidua marker) is shown in yellow. (C) Representative immunofluorescence images of uteri on postpartum day 0.5 or 3.5. Outlined arrowheads indicate GFP-positive luminal epithelial cells. Asterisks indicate placental detachment sites. Brightness/contrast were adjusted independently for each sample. (D) Representative immunofluorescence image of a dam uterus collected after recovery from 4 consecutive pregnancies. (E) Summary schematic of luminal epithelium remodeling around the decidua at implantation sites on gestational day 7.5, represented as longitudinal and transverse views. Single channel outsets are shown as maxIPs, and insets are shown as single Z planes. Inset channel is indicated by box color unless directly noted. Dashed lines indicate bilaminar arches. Outset scale bars, 500 µm; inset scale bars, 50 µm.

## Discussion

Our data indicate that the luminal epithelium plays a pivotal role across numerous endometrial remodeling events in mice. During induced menstruation, we identified prominent bilaminar epithelial arches—largely devoid of *Cxcl15-* lineage cells—that cup the shedding decidual masses along the length of the horn. Our data support a model in which these arches proliferate and extend to envelop the deciduas, enabling concurrent re-epithelialization and shedding without substantial contributions from *Cxcl15-*lineage glandular progenitors. Cell migration may also contribute to the enveloping process.^30,37^ We identified analogous *Cxcl15*-lineage-negative epithelial arches that envelop the decidua during gestation, suggesting that menstruation and gestation share common remodeling mechanisms, potentially mediated by signaling from the decidua or basal endometrium. Whether the expansion of the luminal arch actively facilitates decidual detachment is an exciting possibility for future work. The endometrial epithelium in *Blimp1* knockout mice fails to both encapsulate and expel gestational decidua,^38^ consistent with functional coupling of these processes.

Our data are inconsistent with a model in which glandular progenitors are the primary drivers of murine endometrial re-epithelialization.^14,19^ However, we cannot exclude the possibility of low levels of gland progenitor contributions accumulating across multiple rounds of injury. The discrepancy between our lineage tracing results and those supporting major gland contributions may arise from technical limitations of lineage tracing: even rare ectopic *Cre* driver expression in the highly proliferative luminal epithelium would be amplified to produce labeled epithelial patches that could be interpreted as gland contributions. Rather than serving as the primary source of luminal epithelium under physiological conditions, our data suggest that glandular populations contain facultative progenitor cells, which can be induced to generate functional luminal epithelial cells after extensive chemical ablation of the endometrial surface. In recent work we have reciprocally demonstrated that luminal epithelial cells can be induced to convert into functional glandular epithelium, indicating that lineage plasticity in the mouse endometrial epithelium is bidirectional.^39^

In humans, there is evidence that both glandular and luminal cells contribute to endometrial resurfacing.^7-10,13,37,40^ Human menstrual shedding occurs in a piecemeal manner, with denuded regions interspersed among luminal epithelium which has not yet shed, or has recently regenerated.^6^ Our data from the mouse indicate that persistent luminal epithelium preferentially resurfaces tissue if available, but extensive loss of luminal epithelium evokes gland contributions. If progenitor choice in the human endometrium is governed by similar principles, the relative contributions of luminal and glandular progenitors may vary locally according to the extent of regional luminal epithelium loss. Additionally, the widespread luminal denudation associated with parturition^41,42^ may favor a greater contribution from the glands.

Although the respective contributions of glandular and luminal populations to human endometrial regeneration are challenging to determine definitively, luminal epithelium participation would have important implications for endometrial pathologies. Glandular progenitors have been favored as cells of origin for endometrial cancers and endometriosis,^43-45^ but our data indicating potential progenitor function of the luminal epithelium warrants investigation into whether these cells can also seed tumors or endometriosis lesions.

Whether the mechanisms of luminal epithelium resurfacing identified here extend to other naturally menstruating mammals remains to be determined, but observations from the literature provide tantalizing suggestions. Drawings from the 1950s depicting the *Elephantulus* (elephant shrew) endometrium during menstruation^46^ reveal structures and regional epithelial morphologies that are highly similar to the bilaminar epithelial arches we observe (Figure S4). Interestingly, the only known menstruating rodent, *Acomys* (spiny mouse), has been found to lack endometrial glands altogether,^47^ raising the possibility that luminal epithelium-dominant mechanisms of endometrial re-epithelialization may play a significant role in this species.

Beyond progenitor choice, our findings point to a potential broader principle of endometrial regeneration in which tissue denudation is minimized at any given time. In mice, the coordination of luminal epithelium expansion with tissue detachment provides an elegant solution for tissue elimination at massive scale, while limiting the exposure of sensitive stromal tissues. Piecemeal shedding in human menstruation may represent an alternative approach to achieve the same goal. Because the endometrium programs its own disruption during menstruation and pregnancy, it is uniquely positioned to coordinate extensive repair before tissue excision is complete, which may contribute to its high regenerative capacity.

### Limitations of the Study

This study employs the *Cxcl15*^*iCre*^; *R26*^*nTnG*^ mouse line to track contributions from glandular epithelial populations to the luminal epithelium. We use *Cxcl15* as a marker of gland lineage rather than making claims about its function; of note, *Cxcl15* lacks a human homolog. In principle, emergence of GFP-positive luminal epithelium upon chemical ablation could arise from activation of *Cxcl15* expression, rather than *Cxcl15* lineage descent. However, we detected gland proliferation and epithelial cells emerging from gland openings, consistent with gland-driven re-epithelialization.

The *Cxcl15*^*iCre*^; *R26*^*nTnG*^ lineage tracing tool is highly efficient but does not label every glandular cell. Rare GFP-negative, FOXA2-positive cells were distributed throughout the glands, suggesting that incomplete labeling may reflect stochastic variation in *Cre* recombination rather than the exclusion of a region-specific glandular subpopulation. Non-epithelial cells including fibroblasts, pericytes, and bone marrow-derived cells, have been proposed to generate up to 17% of the epithelium through trans-differentiation.^48-51^ Due to the absence of luminal epithelium-specific tracing tools, we cannot perform a definitive assessment of the relative contributions of luminal progenitors compared to potential non-epithelial sources, although epithelial lineages are not depleted over time in homeostasis, or after multiple pregnancies.^19^ Additionally, luminal epithelium populations are heterogenous,^52,53^ and a progenitor population proposed to reside at the intersection zone between glands and luminal epithelium^20^ may be among the *Cxcl15*-negative cells that participate in luminal resurfacing. Further refining the molecular identities of progenitors among the *Cxcl15*-negative population is an important avenue for future study.

Several aspects of the menstrual remodeling process also remain unresolved. Bilaminar arch structure varies within and between individuals and across menstruation models, and further investigation is required to understand the factors governing their formation and architecture. We speculate that arches initially arise from displacement and compression of pre-existing luminal epithelium by the expanding decidua. We also identified substantial apoptosis in early post-menstrual remodeling and its role in maintaining epithelial lineage balance merits further investigation. Finally, how the luminal epithelium reconnects with the retained gland bases remains an open question.

## Methods

All key resources used in this study are listed in the Key Resources Table

### Animal husbandry

All *Mus musculus* experiments were approved by the Harvard University Institutional Animal Care and Use Committee and the University of Missouri’s Institutional Animal Care and Use Committee and adhered to all relevant ethical regulations and guidelines. All mice were housed with a 14:10 hour light-dark cycle maintained at 21 – 25°C and 30 – 70% humidity, with food (irradiated LabDiet Prolab Isopro RMH 3000 5P75; LabDiet, St. Louis, MO) and water available *ad libitum*. Female mice were group-housed with no more than 5 per cage (7.75” W x 12” L x 6.5” H), with Coarse Grade Sani-Chip Bedding (P.J. Murphy, Ladysmith, WI) and used for experiments between 8 and 21 w of age. When indicated, estrous cycle staging was carried out by performing vaginal lavages with 30 – 100 µL of sterile saline and microscopically assessing the ratios of exfoliated vaginal cornified epithelial cells, nucleated epithelial cells, and leukocytes present in the wet unstained vaginal lavage fluid between 3:00 and 4:00pm.^54^ When indicated, induction of pseudopregnancy (a physiological state of high progesterone) was achieved by pairing females with single vasectomized CD-1 males. The day the copulatory plug was found was designated pseudopregnancy day 0.5. For induction of pregnancy, females were paired with single fertile C57BL/6J (for fertility studies and gestational lineage tracing) or *R26*^*mTmG*^ males (for lineage tracing fetal tissues) and checked for the presence of a copulatory plug every morning and evening until found. The day the copulatory plug was found was designated gestational day 0.5. For fertility testing and postpartum experiments, pregnant dams were checked each morning for new litters starting on gestational day 17.5. Pups were counted when relevant. For postpartum experiments, the day the litter was found was considered postpartum day 0.5. All experiments were performed with at least n = 3 mice per condition.

### Tissue collection

To label proliferating cells *in vivo*, mice received an intraperitoneal injection of 25 µg/g EdU 1 h prior to tissue harvest. Animals were either euthanized by 5 min exposure to CO_2_ or by cervical dislocation. Uterine horns were promptly excised with ovaries, cervix and mesometrium attached. Horns were laid flat on filter paper, briefly imaged in a 10” x 10” LED lightbox to document gross morphology, then fixed in ice cold 4% paraformaldehyde overnight at 4°C with gentle agitation. Samples were then washed 3 times, 15 min – 2 h each in PBS, and trimmed of any excess fat or mesometrium as needed, before storage at 4◦C in 0.1% azide in PBS until use.

### *Cxcl15*^*iCre*^; *R26*^*nTnG*^ lineage tracing experiments

Uteri from *Cxcl15*^*iCre*^ *R26*^*nTnG*^ females were collected at the following time points for lineage tracing experiments:

- Homeostasis (including 9 and 21 week old samples): afternoon of pseudopregnancy day 4.5, unless a specific estrous cycle stage is indicated.
- Estrous cycle stages: afternoon of lavage fluid collection, between 3:00pm and 5:00pm. All mice were between 9 and 11 weeks of age.
- Chemical ablation: polidocanol injection was performed when animals were in estrus, and horns were collected for further analysis 2 h, 72 h, 14 d, or 28 d later.
- Oil menstruation: either (1) the day of their first robust positive guaiac test, (2) the day after bleeding cessation, defined as the day after the first negative guaiac test following an initial positive test, or (3) 10 d after the last estradiol and progesterone injection.
- Gestation: gestational day 7.5 or 8.5 between 12:00pm and 4:00pm.
- Postpartum: postpartum day 0.5 or 3.5 between 12:00pm and 4:00pm.

### Luminal epithelium chemical ablation

C57BL/6 or *Cxcl15*^*iCre*^; *R26*^*nTnG*^ females in the estrus stage of the hormonal cycle were identified by the ratio of leukocytes, nucleated epithelial cells, and cornified epithelial cells present in vaginal lavage fluid. Polidocanol foam was generated by combining 4 parts air and 1 part 5% w/v polidocanol in syringes attached with a luer lock adapter and passing the mixture through the adapter 60 times each way. Polidocanol foam or saline (for negative controls) was injected into the uterine lumen from the cervical third of the horn using a 32G needle until visual confirmation that it filled the entire horn, then the process was repeated for the contralateral horn. Any foam that leaked out of the injection site was thoroughly flushed away with sterile saline. The foam or saline was allowed to sit in the uterine lumen for 15 min to facilitate full ablation of the luminal epithelium. During this time, the abdominal cavity was covered with sterile gauze to limit exposure to the environment, and routinely irrigated with warmed sterile saline to keep the tissue hydrated. After 15 min, the foam was flushed out of the lumen by injecting 250 µL sterile saline per horn using a 32G needle to limit further ablation. Residual polidocanol foam was thoroughly flushed from the outside of the uterus, abdominal fat pads and skin around the incision site with sterile saline immediately after polidocanol injection and intrauterine saline flushing. The uterus was then returned to the abdominal cavity and the incision was closed. Uterine horns were collected at the indicated time points or animals were allowed to recover for 4 days for fertility testing (see Figure 2E for experimental timeline). All mice used for this assessment exhibited copulatory plugs within 9 days of polidocanol treatment.

### Oil induction of menstruation in ovariectomized mice

Cxcl15^iCre^; R26^nTnG^ females were bilaterally ovariectomized, the*Cxcl15*^*iCre*^; *R26*^*nTnG*^ females were bilaterally ovariectomized, then allowed to recover for 7 – 14 d (see Figure S2A for an experimental timeline). Beginning on day 0, mice received 3 consecutive days of subcutaneous estradiol (100 ng in sesame oil), then were allowed to rest for 2 days. Mice then received daily subcutaneous injections of 6.7 ng of estradiol and 1 mg progesterone in sesame oil from days 5 – 11. To induce decidualization, mice received an intrauterine oil injection on the evening of day 7. Each horn received 50 µL of sesame oil or sterile saline (for negative controls) using a 31G insulin syringe inserted into the third of the horn proximal to the cervix. To assess bleeding, daily vaginal lavages were collected in 30 – 100 µL sterile saline from mice beginning the day after intrauterine oil injection. Lavage fluid was tested for the presence of red blood cells using guaiac feccal occult blood tests. Animals were harvested at one of three timepoints as indicated in Figure S2A: (1) the day of their first robust positive guaiac test, (2) the day after bleeding cessation, defined as the day after the first negative guaiac test following an initial positive test, or (3) 10 d after the last estradiol and progesterone injection. Animals that exhibited positive guaiac tests before the last progesterone injection were omitted from the study.

### Chemogenetic induction of menstruation in X-Mens mice

We used the previously described method of chemogenetic induction of decidualization and menstruation in X-Mens mice.^34^ Pseudopregnancy was induced in non-ovariectomized *Amhr2*^*Cre*^; *R26*^*GsD*^ females to raise progesterone levels (see Figure S2D for an experimental timeline). On the evening of pseudopregnancy day 3.5, animals received drinking water with 48 μg/mL clozapine-N-oxide dihydrochloride (CNO) after 4:00 pm. On day 4.5, animals were given 5 consecutive intraperitoneal injections of a DREADD agonist cocktail of 24 μg/mL CNO, 125 μg/mL deschloroclozapine dihydrochloride, and 200 μg/mL compound 21 dihydrochloride in physiological saline, spaced 2 h apart. The CNO drinking water was replaced with regular drinking water by 11:00 am the following day. In this model, DREADD activation under a high progesterone state induces decidualization. As the pseudopregnancy resolves, progesterone levels will naturally decline, which triggers decidual shedding. To assess shedding and bleeding, daily vaginal lavages were collected in 30 – 100 µL sterile saline from mice beginning the day after DREADD agonist cocktail administration. Lavage fluid was tested for the presence of red blood cells using guaiac fecal occult blood tests and animals were harvested on the day of their first robust positive guaiac test.

### Consecutive pregnancies

*Cxcl15*^*iCre*^ *R26*^*nTnG*^ females (6 – 8 weeks of age) were housed individually and continuously with a fertile *Cxcl15*^*iCre*^ *R26*^*nTnG*^ male to allow uninterrupted breeding. Females were maintained with the male until four litters were produced, after which the male was removed. The fourth litter was weaned at postnatal day 21, and uteri were collected from the dam during the subsequent diestrus. The tissue was harvested and fixed in 4% paraformaldehyde in PBS for 60 min at 4 °C, followed by cryoprotection by overnight immersion in 15% sucrose in PBS at 4 °C. Tissue was embedded in optimal cutting temperature compound (OCT) and cryosectioned at 7 – 10 μm. Sections were mounted on Superfrost Plus slides, counterstained with 2 μg/mL Hoechst 33342, and coverslipped with ProLong™ Diamond Antifade Mountant. Images were acquired using a Leica DM6 B upright microscope equipped with a Leica K8 camera and Leica Application Suite X (LAS X) software.

### Tissue cryosectioning and immunostaining

Following overnight fixation, individual uterine horns were transferred to 30% sucrose in PBS and incubated overnight at 4 °C for cryoprotection. The two uterine horns were separated at the cervix, then embedded in OCT compound, snap frozen, and stored at –80 °C until use. For transverse sections, horns were bisected transversely prior to embedding. OCT blocks were equilibrated in the cryostat chamber at –19 to –21 °C for a minimum of 30 min prior to sectioning. Horns were sectioned on a Leica CM 1860 cryostat either longitudinally at 50 µm thickness, or transversely at 12 μm thickness starting from the medial (cut) portion of the horn, and working toward the distal (cervical or ovarian) ends. Sections were collected onto positively charged glass slides and stored at –80 °C.

For immunofluorescence, section perimeters were outlined with a hydrophobic barrier pen. The following buffers were used:

- Blocking buffer: Either (1) PBS containing 5% normal donkey serum, 0.5% BSA, 0.1% Triton X-100, and 0.1% sodium deoxycholate or (2) 5% normal donkey serum, 2% BSA, and 0.1% - 0.3% Triton X-100 in PBS
- PBT: Either (1) 0.1% Tween-20 in PBS or (2) 0.1% - 0.3% TritonX-100 in PBS

Slides were optionally post-fixed in 4% formaldehyde in PBS for 5 minutes at room temperature, then washed three times in (PBT), 15 min per wash. Sections were incubated for 30 – 60 min at room temperature in blocking buffer. Sections were incubated with primary antibodies diluted in blocking buffer overnight at 4 °C in a humidified chamber, then washed three times in PBT. Sections were then incubated in blocking buffer containing secondary antibodies and DAPI (1 µg/mL) overnight at 4 °C or 1 h at room temperature in a humidified chamber. Slides were washed three times in PBT or PBS, then coverslipped with ProLong Gold Antifade mounting medium and allowed to cure for at least 24 h at room temperature. All antibodies used are listed in the key resources table.

### Thick section tissue clearing, EdU staining, and immunostaining

Single uterine horns were mounted in 2-5% low gelling temSingle uterine horns were mounted in 2-5% low gelling temperature agarose, then sectioned with a vibratome or compresstome at 200 µm thickness and stored in PBS with 0.1% azide at 4° C until use. All steps were carried out at room temperature, unless noted otherwise. Sections were first incubated at 37°C in permeabilization solution 1 (0.2% TritonX-100, 10% DMSO, 0.2% Deoxycholate, 20 mM EDTA in PBS, pH 8.0) for 24 h, then transferred to permeabilization solution 2 (0.2% Triton X-100, 10% DMSO, 100 mM glycine in PBS) for 1 h at 37°C. X-Mens samples containing endogenous fluorescent proteins required an additional bleaching step, in which sections were incubated in 10% hydrogen peroxide in 0.05 M phosphate buffer at 37°C for 3 h, then quickly washed twice in PTwH (0.2% Tween-20, 10 µg/mL heparin in PBS), followed by three longer washes in PTwH, 10 min each. For EdU staining, sections were quickly washed twice in 3% BSA in PBS, then incubated for 1 h in a freshly made reaction cocktail (2 mM Copper sulfate pentahydrate, 8 µM sulfo-cyanine3 azide and freshly dissolved 113 mM ascorbic acid in PBS, pH 7.4) and then quickly washed twice with 3% BSA before proceeding to blocking. Sections were blocked in 0.2% Triton X-100, 10% DMSO, 5% Normal Donkey Serum in PBS for 1 h, then incubated with primary antibodies diluted in a staining solution of 0.2% Tween-20, 10 µg/mL heparin, 5% DMSO, and 5% Normal Donkey Serum in PBS at 4°C overnight. Sections were then quickly washed twice with PTwH, then three times with PTwH, 10 min each, then allowed to incubate for 3 h with the appropriate donkey secondary antibodies. Staining solutions contained DAPI diluted 1:1000 to label nuclei. Sections were then washed in PTwH twice for a few seconds each, followed by three washes of 10 min each. Next, sections were optically cleared to help facilitate imaging through the full thickness as follows: after removing as much liquid as possible, sections were incubated in an EZ index solution of 80% Nycodenz, 42% Urea and 0.025% sodium azide in 0.2 M, pH 7.4 phosphate buffer for at least 1 h. Sections were mounted in EZ index on slides fit with 200 µm thick spacers and imaged within 4 d of mounting. All antibodies used are listed in the key resources table.

### Whole horn tissue clearing and immunostaining

Single uterine horns were cut in half transversely to facilitate reagent penetration into the sample. Each horn half was processed in parallel through the following steps. The following protocol was used for whole mount staining of both estrus and oil menstruation samples but the THEED and CHAPS steps were excluded for the estrus samples. Unless otherwise noted, horns were incubated in 5 mL of all solutions at 37◦C with vigorous shaking. To elute heme, horns were first incubated for 3 d in 25% w/v THEED (N,N,N’,N’-Tetrakis(2-hydroxypropyl) ethylenediamine) prepared in distilled water; THEED was replaced with a fresh solution each day. THEED was removed by washing with PBS for 30 min three times at room temperature. Next, pigment was removed by incubating samples in 10% hydrogen peroxide in 0.05 M phosphate buffer for 24 h. Then the peroxide solution was replaced with permeabilization solution 1 (0.2% TritonX-100, 10% DMSO, 0.2% Deoxycholate, 20 mM EDTA in PBS, pH 8.0) and allowed to incubate. After 24 h, the samples were placed in permeabilization solution 2 (0.2% Triton X-100, 10% DMSO, 100 mM glycine in PBS) for 24 h. For additional permeabilization, samples were then transferred to 10% w/v CHAPS (3-((3-cholamidopropyl) dimethylammonio)- 1-propanesulfonate) freshly prepared in PBS for 48 h. CHAPS was removed by washing with PBS three times, 30 min each, at room temperature. Samples were next blocked in 0.2% Triton X-100, 10% DMSO, 5% Normal Donkey Serum in PBS for 24 h. Next, samples were incubated with primary antibodies diluted in a staining solution of 0.2% Tween-20, 10 µg/mL heparin, 5% DMSO, and 5% Normal Donkey Serum in PBS for 7 d. Primary antibody solution was removed by washing five times, 1 h each, in PTwH (0.2% Tween-20, 10 µg/mL heparin in PBS). Samples were then incubated with secondary antibodies diluted in a staining solution of 0.2% Tween-20, 10 µg/mL heparin, 5% DMSO, and 5% Normal Donkey Serum in PBS for 7 d. Secondary antibody solution was removed by washing two times, 1 h each, in PTwH, followed by two 1 h washes in PBS, and two 1 h washes in distilled water. To control orientation for clearing and imaging, samples were mounted in 1% low melting temperature agarose prepared in distilled water and immediately transferred to 20% methanol diluted in water for at least 2 h without agitation to begin clearing. All clearing steps were performed at room temperature, in at least 25 mL of solution per sample. Samples were then transferred to 40% methanol overnight with gentle agitation. Clearing was then completed by moving samples through the following sequence of solutions with gentle agitation: 60% methanol for 3 h, 80% methanol for 2 h, 100% methanol for 1 h, 66% dichloromethane diluted in methanol for 3 h, and finally, two 15 min rinses in 100% dichloromethane. Finally, to match the refractive indices of the tissue sample and the imaging medium, samples were transferred to fresh dibenzyl ether and left for at least 12 h without agitation before imaging in dibenzyl ether.

### Imaging and image processing

To show multiple gland-luminal epithelium junctions, immunofluorescence images of 200 µm-thick transverse or 50 µm-thick longitudinal sections are shown as maximum intensity projections (maxIPs), unless noted otherwise in the figure legend. Immunostained uterine sections and whole mount estrus uterine horns were imaged on a Yokogawa CSU-W1 spinning disc confocal system fit to an inverted Nikon Ti-2E microscope. All images of uterine sections were acquired using a CFI60 Plan Apochromat Lambda 20x/0.75 NA objective. 200 µm-thick transverse and 50 µm-thick longitudinal sections were imaged with a 5 µm Z-step size, and 12 µm thick transverse sections were imaged with a 0.9 µm Z-step size. All images of estrus whole mount horns were acquired using a CFI60 Plan Apochromat Lambda D 10x/0.45 NA 0.45 objective with a 5 µm Z-step size. Large images were acquired using Nikon Elements JOBS module. Stitched images and maximum intensity projections were generated using Nikon Elements General Analysis 3 module. PNGs were generated using Fiji/ImageJ. Immunostained whole mount oil-induced menstruation horns were imaged on a Zeiss Lightsheet.7 microscope using an EC Plan-Neofluar 5x/0.16 NA (nD=1.33-1.58) objective lens and a 5µm Z-step size. All representations of whole mount imaging data including 3D renderings, maxIPs and movies were generated using Zeiss Arivis Pro software.

### Quantification and statistical analysis

For all experiments, at least three independent replicates were examined. For each transversely sectioned replicate, three sections collected from the cervix-proximal, middle, and ovary-proximal thirds of each uterine horn were examined. Data are expressed as mean +/-standard deviation. All section images were analyzed in Fiji/ImageJ.

For quantification of the proportion of GFP-positive glandular epithelium in *Cxcl15*^*iCre*^; *R26*^*nTnG*^ samples, 200 µm-thick sections were immunostained for FOXA2 and GFP. For each section, three equally spaced Z-planes were selected for analysis. FOXA2-positive cells were first manually annotated using the multi-point tool. After annotating all FOXA2-positive cells on a given section, the GFP channel was toggled on and the previously marked FOXA2-positive cells were re-inspected; cells were scored as double-positive (FOXA2-positive/GFP-positive) when GFP signal overlapped with the same cell. Any candidate cell with a mean pixel intensity less than 300 (arbitrary units) was deemed too faint to accurately classify and excluded from analysis.

For quantification of the proportion of GFP-positive luminal epithelium in *Cxcl15*^*iCre*^; *R26*^*nTnG*^ samples, three 12 µm-thick cryosections per uterine horn (spaced at least 300 µm apart) were immunostained for GFP using an AF-647 secondary antibody to maximize the ratio of signal to noise. Luminal epithelial cells were defined as the layer of nuclei in contact with the lumen, as shown with the dashed lines in Figure 1F insets. A gap between these nuclei and the nuclei of the underlying stroma facilitates identification of this layer. This conservative definition of luminal epithelium includes the junctions between glandular epithelium and the epithelium lining the lumen and thus includes a small number of GFP-positive/FOXA2-positive cells. For homeostasis (pseudopregnancy) and oil menstruation conditions, the mean number of luminal epithelial nuclei per µm was determined to be 0.475 by quantifying the number of nuclei spanning a 40 µm long luminal segment across randomly selected regions; 21 regions in total were measured across 7 sections (3 regions per section). For estrous cycle stage comparisons, the mean number of luminal epithelial nuclei per µm was determined independently for each cycle stage to account for varying nuclear morphologies across the estrous cycle; for each cycle stage, 9 regions were measured across 3 sections (3 regions per section). The number of nuclei per µm was 0.37 in diestrus, 0.43 in proestrus, 0.53 in estrus, and 0.34 in metestrus. The perimeter of the uterine lumen was then traced in each section using the polygon selection tool in Fiji, and the total luminal epithelial cell number was estimated by multiplying the measured perimeter by the mean number of nuclei per µm for the corresponding condition. GFP-positive cells were counted manually. All quantification data are provided in Table S1.

For bilaminar arch frequency quantification, entire uterine horns were serially sectioned at a 200 µm-thickness. Every other section was immunostained for EPCAM and examined for the presence of continuous EPCAM-positive cells (sheets) around at least a portion of the periphery of the decidua. Damaged sections were excluded from analysis.

Statistical analyses were completed using Prism 11 software. Comparisons between the proportion of mice that successfully carried pregnancies to term following either saline or polidocanol treatment were conducted using Fisher’s exact test. Comparisons between the ratio of GFP-positive luminal epithelium across each of the four estrous cycle stages were performed using a one-way ANOVA, followed by Tukey’s multiple comparisons test to identify which cycle stages differed. All other comparisons between the ratios of GFP-positive luminal epithelium or litter size across two conditions were performed using Welch’s unpaired t-tests.

## Resource Availability

Requests for further information and resources should be directed to and will be fulfilled by the lead contact, Kara McKinley. This study did not generate new unique reagents or code.

## Acknowledgements

We thank Iain Cheeseman, Ya-Chieh Hsu, Pradeep Tanwar and McKinley and Kelleher lab members for feedback. We thank Kathleen Pritchett-Corning, Eny Maldonado, and the Harvard Office of Animal Resources for animal care, the Harvard Center for Biological Imaging (HCBI), and the Histology Core at the Harvard Stem Cell and Regenerative Biology Department for critical resources, and Timothy Fenn for administrative support. We extend our deepest gratitude to our families and communities. This work was supported by the National Institutes of Health (DP2HD111708 and R00HD101021 to KLM, R01HD112315 to AM Kelleher, R37HD114609 to TES), New York Stem Cell Foundation (KLM), Howard Hughes Medical Institute (KLM), Charles H. Hood Foundation (KLM), David and Lucile Packard Foundation (KLM), Searle Scholars Program (KLM), Smith Family Foundation (KLM) and Damon Runyon Cancer Research Foundation (KLM). KLM is a New York Stem Cell Foundation Robertson Stem Cell Investigator. This material is based on work supported by the National Science Foundation Graduate Research Fellowship Program to CJA under grant number DGE 2140743. CJA was additionally supported by the Simmons Awards and the Vranos Family Foundation. ÇÇ was supported by the Soozin White Research Fund. FSC was supported by the Long-term Fellowship from the Human Frontiers Science Program and the Postdoc.Mobility fellowship from the Swiss National Science Foundation.

## Author Contributions

Conceptualization: CJA, KLM Methodology: CJA, JLRG, TDS, PLM

Visualization: CJA, ÇÇ, PLM

Investigation: CJA, JLRG, EMW, KCL, ÇÇ, TDS, PLM, SGB,

BDM, AM Kaage, DM, AB, AEG

Funding acquisition: CJA, KLM

Project administration: CJA, ÇÇ, AM

Kelleher, KLM

Supervision: CJA, AM

Kelleher, KLM

Writing – Original Draft: CJA

Writing – Review and Editing: CJA, JLRG, EMW, KCL, ÇÇ, TDS, PLM, SGB, BDM, AM

Kaage, DM, AB, AEG, FSC, TES,

AM Kelleher, KLM

Resources – TES, AM Kelleher

Formal analysis: CJA, EMW, KCL, ÇÇ, PLM, SGB, BDM, AB, AEG, FSC

Data curation: CJA

## Declaration of Generative AI and AI-assisted technologies in the writing process

During the preparation of this work the authors used Claude and ChatGPT within the Harvard AI Sandbox to provide suggested edits to improve clarity. After using these tools, the authors reviewed and edited the content as needed and take full responsibility for the content of the publication.

