## Supplementary materials for "Luminal epithelium remodeling underlies endometrial regeneration during menstruation and pregnancy"

This file includes:
Figures S1, S2, S3 and S4
Table S1
Video S1 and S2 legends
Key Resources Table

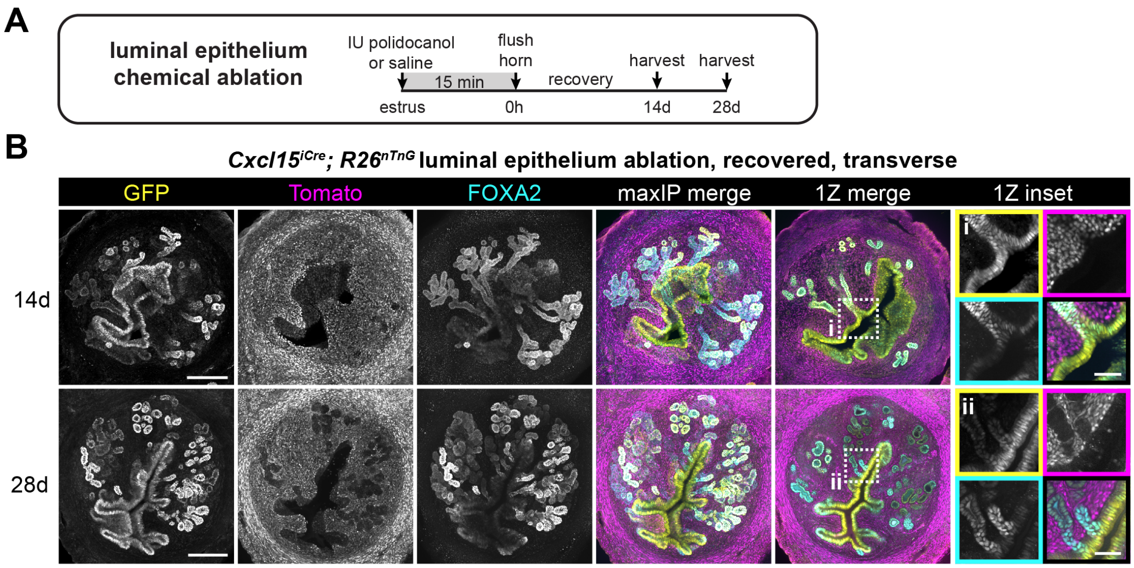


**Figure S1. Cxcl15 lineage luminal epithelium persists through long term recovery from luminal epithelium ablation, related to Figure 2.**

(A) Experimental timeline for luminal epithelium chemical ablation.

(B) Representative immunofluorescence images of uteri collected as indicated in A.

For all images, single channel outsets are shown are as maxIPs. Inset channel is indicated by box color. Outset scale bars, 200 μm; inset scale bars, 50 μm.

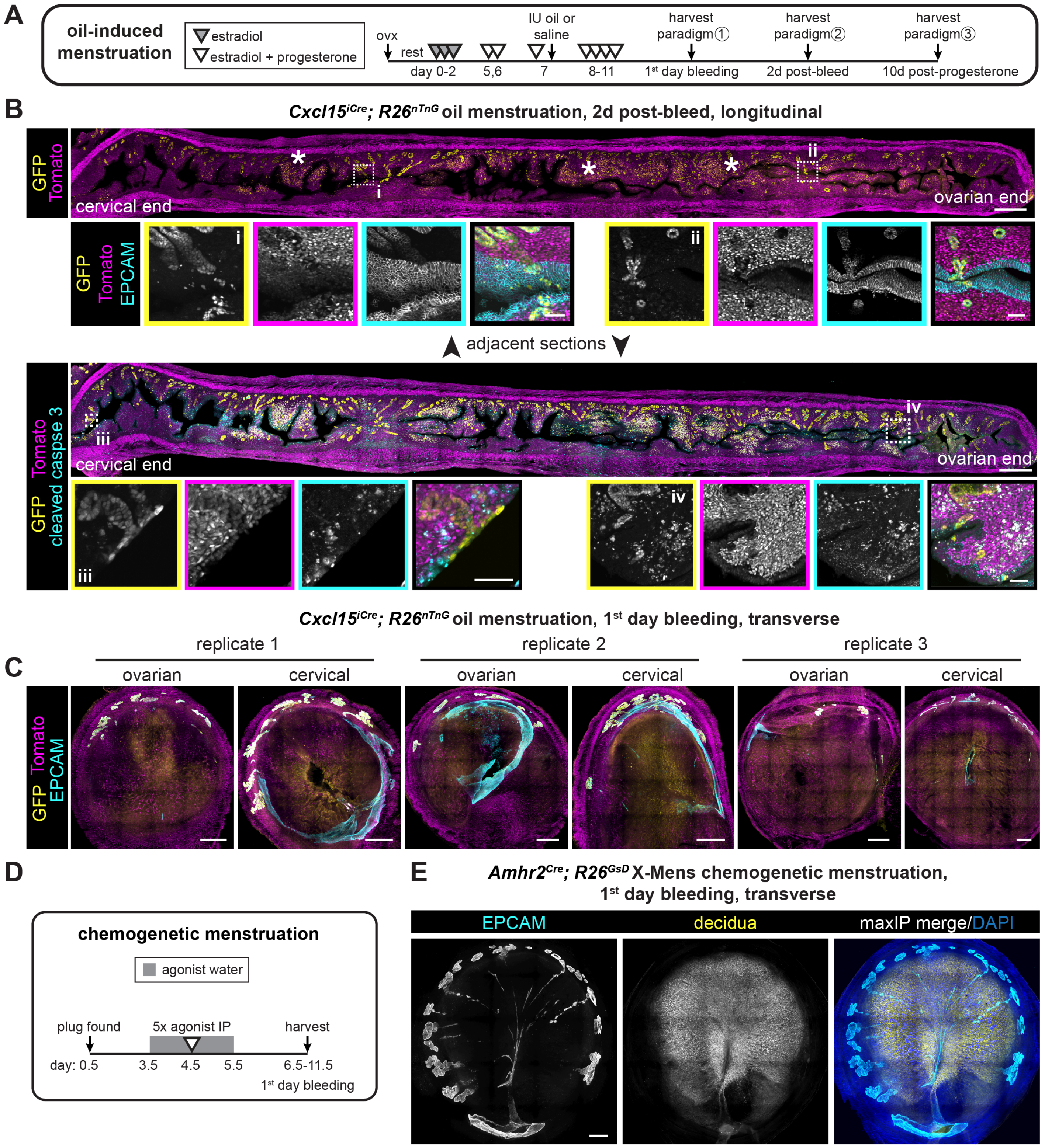


**Figure S2. Morphogenetic changes to luminal epithelium throughout menstruation, related to Figure 3.**

(A) Experimental timeline for oil induction of menstruation in ovariectomized mice.

(B) Representative adjacent uterine sections collected two days after cessation of oil-induced menstrual bleeding and indicated in panel A, paradigm 2. Asterisks indicate examples of GFP-positive cells in the stromal layer, which frequently co-localize with the apoptosis marker, cleaved caspase 3.

(C) Additional examples of immunofluorescence images of uterine sections collected on the first day of oil-induced menstrual bleeding. Two regions per horn are shown (ovary-proximal and cervix-proximal). A third, medial region from replicate 2 is shown in Figure 3F. Brightness/contrast were adjusted independently for each sample.

(D) Experimental timeline for chemogenetic induction of menstruation in X-Mens mice.

(E) Representative immunofluorescence images of an X-Mens uterus exhibiting a bilaminar arch structure, collected as indicated in D. 27% of serial sections examined exhibited a bilaminar arch, and all sections examined contained persistent luminal epithelium.

All images are shown as maxIPs. Inset channel is indicated by box color. Outset scale bars, 500 μm; inset scale bars, 50 μm. IP, intraperitoneal injection.

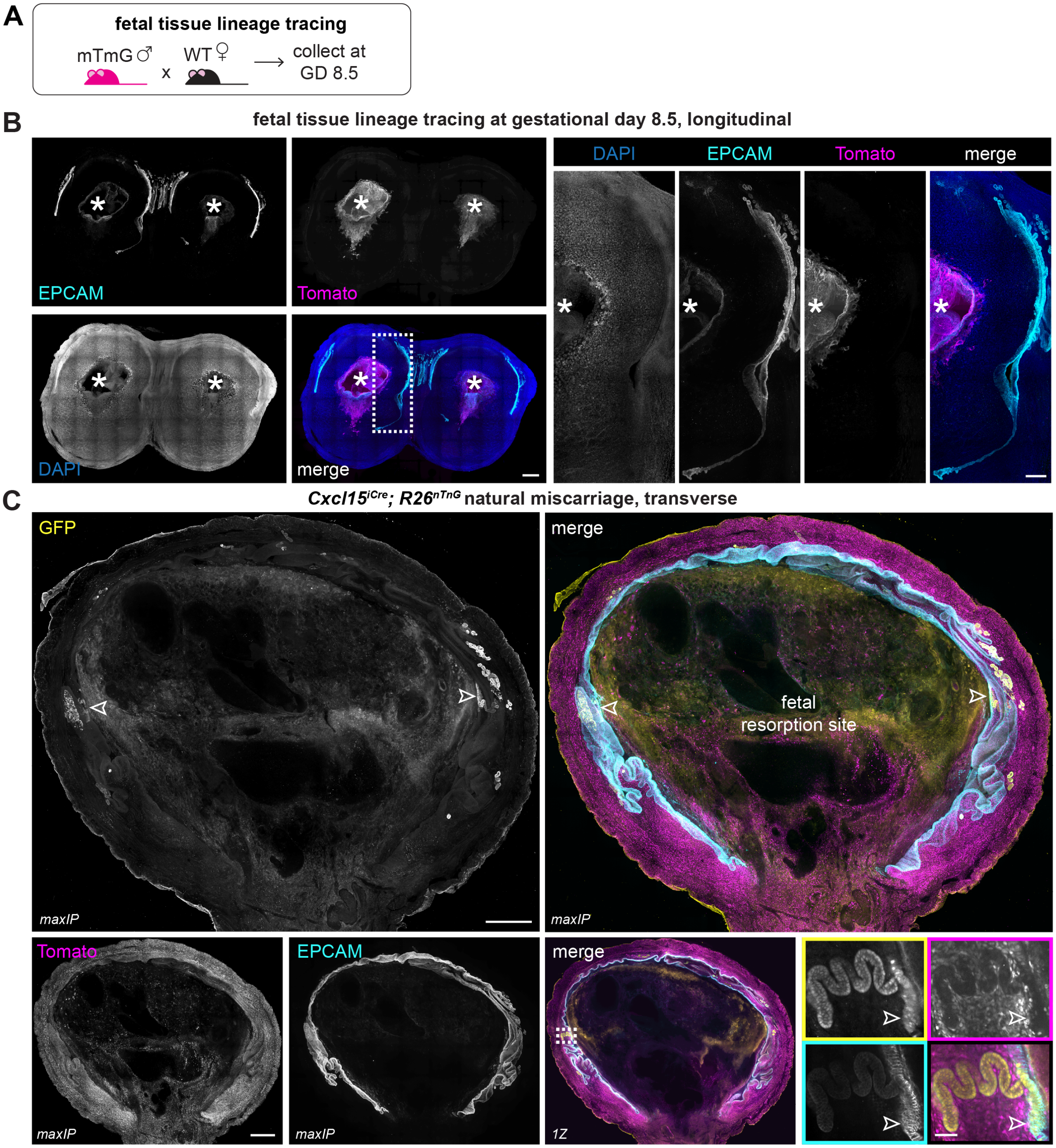


**Figure S3. *Cxcl15* lineage independent luminal epithelium contributions to gestational remodeling and fetal resorption, related to Figure 4.**

(A) Experimental paradigm for lineage tracing fetal-derived tissue.

(B) Representative immunofluorescence maxIP images of two adjacent longitudinally sectioned implantation sites, collected as indicated in A. Asterisks indicate embryos. Outset scale bars, 500 µm; inset scale bars, 200 µm.

(C) Single immunofluorescence image of a naturally arising fetal-resorption site collected on the day of parturition. Outlined arrowheads indicate rare GFP-positive luminal epithelial cells. Outset scale bars, 500 µm; inset scale bars, 50 µm.

Insets are shown as single Z planes. Inset channel is indicated by box color unless directly noted. GD, gestational day; WT, wildtype.

**menstruating elephant shrew uterus, transverse**

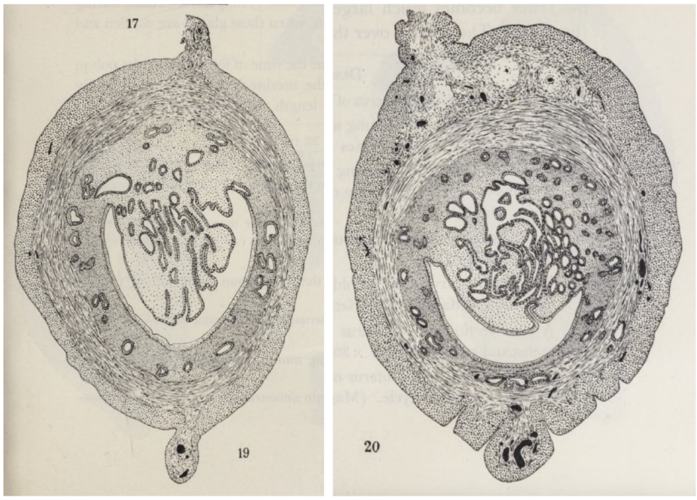


young animal aged animal

**Figure S4. Epithelial morphology in menstruating elephant shrew, related to Figure 3.**

Drawings of uterine morphology during menstruation in young and aged elephant shrews from C.J. Van der Horst's 1954 manuscript, *Elephantulus Going into Anoestrus; Menstruation and Abortion.*[46] Reproduced with permission from The Royal Society Publishing.

| Mouse ID | Avg number of GFP+ luminal epithelium nuclei per section | Avg number of luminal epithelium nuclei per section | Percent GFP+ luminal epithelium nuclei per section | Avg percent GFP+ luminal epithelium nuclei across condition ± std dev |
|---|---|---|---|---|
| **Homeostasis, diestrus** | | | | |
| W502 | 7.00 | 1656.40 | 0.42 | 0.33 ± 0.29 |
| X403 | 0.00 | 672.33 | 0.00 | |
| X941 | 4.67 | 835.09 | 0.56 | |
| **Homeostasis, proestrus** | | | | |
| X942 | 6.33 | 1776.90 | 0.36 | 0.77 ± 0.52 |
| W694 | 22.67 | 3774.55 | 0.60 | |
| W627 | 38.33 | 2841.20 | 1.35 | |
| **Homeostasis, estrus** | | | | |
| X940 | 3.33 | 1940.21 | 0.17 | 0.53 ± 0.33 |
| W631 | 33.00 | 4039.40 | 0.82 | |
| W633 | 15.33 | 2538.58 | 0.60 | |
| **Homeostasis, metestrus** | | | | |
| W510 | 16.0 | 1199.77 | 1.33 | 1.38 ± 0.37 |
| W999 | 24.33 | 2358.56 | 1.03 | |
| W507 | 23.00 | 1292.74 | 1.78 | |
| **Homeostasis, 9w of age** | | | | |
| L524 | 0.67 | 6,052.14 | 0.01 | 0.09 ± 0.09 |
| L525 | 12.30 | 6,484.77 | 0.19 | |
| L527 | 2.67 | 4,554.60 | 0.06 | |
| **Homeostasis, 21w of age** | | | | |
| L522 | 7.00 | 3,681.19 | 0.20 | 0.19 ± 0.01 |
| L526 | 6.00 | 3,585.47 | 0.17 | |
| L874 | 9.50 | 4,271.05 | 0.19 | |
| **Oil-induced menstruation, control** | | | | |
| U317 | 8.50 | 8,164.38 | 0.10 | 0.06 ± 0.05 |
| U492 | 4.00 | 4,360.32 | 0.09 | |
| Q123 | 0.00 | 2,512.91 | 0.00 | |
| Q124 | 1.33 | 3,143.17 | 0.05 | |
| **Oil-induced menstruation, menstruated** | | | | |
| Q643 | 4.33 | 3,328.54 | 0.13 | 0.27 ± 0.14 |
| Q644 | 3.00 | 2,451.30 | 0.13 | |
| U315 | 21.33 | 7,592.21 | 0.31 | |
| U817 | 7.67 | 3,439.26 | 0.26 | |
| U818 | 15.33 | 2,849.64 | 0.55 | |
| U819 | 8.00 | 1,856.68 | 0.32 | |
| U820 | 5.00 | 1,744.84 | 0.28 | |
| V276 | 11.33 | 7,644.70 | 0.15 | |

**Table S1. Percent GFP-positive luminal epithelium across homeostasis and menstruation, related to Figures 1 and 3.**

Quantification of the proportion of luminal epithelial nuclei expressing GFP in homeostatic conditions, each estrous cycle stage, and after recovery from oil-induced menstruation in *Cxcl15*$^{iCre}$*; R26*$^{nTnG}$ mice. Values for each mouse are averaged across three cryosections.

**Video S1. Continuous epithelial architecture across decidual masses during oil-induced menstrual bleeding, related to Figure 3C.**

Video showing multiple views of the sample in Figure 3C. E-cadherin staining shown in cyan. Video is scaled with ɣ adjustment and adjusted to maximize opacity.

**Video S2. Discontinuous epithelial architecture across decidual masses during oil-induced menstrual bleeding, related to Figure 3D.**

Video showing multiple views of the cervical half of Figure 3D. E-cadherin staining shown in cyan. Video is scaled with ɣ adjustment and adjusted to maximize opacity.

### Key Resources Table

See main text for reference list

| REAGENT or RESOURCE | SOURCE | IDENTIFIER |
|---|---|---|
| **Antibodies** | | |
| Rabbit polyclonal anti-cleaved caspase 3; 1:500 | Cell Signaling Technologies | Cat# 9661-S; RRID: AB_2341188 |
| Chicken polyclonal anti-GFP; 1:1000 | Abcam | Cat# ab13970; RRID: AB_300798 |
| Goat polyclonal anti-P-cadherin; 1:200 | R&D Systems | Cat# AF761; RRID: AB_355581 |
| Goat polyclonal anti-TdTomato; 1:1000 | SICGEN | Cat# AB8181; RRID: AB_2722750 |
| Rabbit monoclonal anti-FoxA2/HNF3 (clone D56D6); 1:200 | Cell Signaling Technologies | Cat# 8186; RRID: AB_10891055 |
| Rat monoclonal anti-CD326/EpCAM (clone G8.8); 1:1000 | BioLegend | Cat# 118202; RRID: AB_1089027 |
| Rat monoclonal anti-Cytokeratin 8 (clone TROMA-1); 0.42 µg/mL final concentration | Developmental Studies Hybridoma Bank (Brûlet et al.[59]) | Cat# TROMA-1 |
| Rabbit monoclonal anti- E-cadherin (clone 24-E10); 1:1000 | Cell Signaling Technologies | Cat# 3195S; RRID: AB_2291471 |
| Donkey anti-Chicken AF488; 1:1000 | Thermo Fisher Scientific | Cat# A78948; RRID: AB_2921070 |
| Donkey anti-Goat AF488; 1:1000 | Thermo Fisher Scientific | Cat# A11055; RRID: AB_2534102 |
| Donkey anti-Goat AF555; 1:1000 | Thermo Fisher Scientific | Cat# A32816; RRID: AB_2762839 |
| Donkey anti-Goat AF647; 1:1000 | Thermo Fisher Scientific | Cat# A21447; RRID: AB_2535864 |
| Donkey anti-Rabbit AF555; 1:1000 | Thermo Fisher Scientific | Cat# A31572; RRID: AB_162543 |
| Donkey anti-Rabbit AF647; 1:1000 | Thermo Fisher Scientific | Cat# A31573; RRID: AB_2536183 |
| Donkey anti-Rat AF405; 1:500 | Thermo Fisher Scientific | Cat# A48268; RRID: AB_2890549 |
| Donkey anti-Rat AF488; 1:1000 | Thermo Fisher Scientific | Cat# A21208; RRID: AB_2535794 |
| Donkey anti-Rat AF647; 1:1000 | Thermo Fisher Scientific | Cat# A78947; RRID: AB_2910635 |
| **Chemicals, peptides and recombinant proteins** | | |
| Agarose, low gelling temperature | Sigma-Aldrich | Cat# A9414 |
| Ascorbic acid | Sigma-Aldrich | Cat# A4544 |
| BSA (bovine serum albumin) | Sigma-Aldrich | Cat# 10775835001 |
| CHAPS Detergent (3-((3-cholamidopropyl) dimethylammonio)-1-propanesulfonate) | Thermo Fisher Scientific | Cat# 28300 |
| CNO (clozapine-N-oxide dihydrochloride) | MedChemExpress | Cat# HY-17366A |
| Compound 21 dihydrochloride | MedChemExpress | Cat# HY-100234A |
| $CuSO_4$ $5H_2O$ (copper sulfate pentahydrate) | Sigma-Aldrich | Cat# 12849 |
| DAPI | Thermo Fisher Scientific | Cat# 62248 |
| DCZ (deschloroclozapine dihydrochloride) | MedChemExpress | Cat# HY-42110A |
| DMSO (dimethyl sulfoxide) | Tokyo Chemical Industry | Cat# D5293 |
| Dibenzyl ether | Thermo Fisher Scientific | Cat# 148400025 |
| Dichloromethane | Sigma-Aldrich | Cat# 27099 |
| EDTA (0.5 M, pH 8.0) | Thermo Fisher Scientific | Cat# 15575020 |
| EdU (5-Ethynyl-2’-deoxyuridine) | TargetMol | Cat# T17341 |
| 17β-Estradiol | Sigma-Aldrich | Cat# E8875 |

| | | |
|---|---|---|
| Ethanol, 200 proof | Decon Labs | Cat# V1016TP |
| Ethiqa XR (extended-release buprenorphine) | Fidelis Animal Health | Cat# NDC 86084-100-30 |
| Glycine | Sigma-Aldrich | Cat# G8898 |
| Heparin | Sigma-Aldrich | Cat# H3393 |
| Hoechst 33342 | Invitrogen | Cat# H3570 |
| Isoflurane | Akorn Inc | Cat# 07-894-6668 |
| Low gelling temperature agarose | Sigma-Aldrich | Cat# A9414 |
| Methanol | Avantor | Cat# 9070-03 |
| Mifepristone | MedChemExpress | Cat# HY-13683 |
| Normal Donkey Serum | Neuromics | Cat# SER004 |
| Nycodenz | Serumwerk | Cat# 18003 |
| Paraformaldehyde, 32% | Electron Microscopy Sciences | Cat# 15714-S |
| PBS with 0.1% azide | Santa Cruz | Cat# sc-296028 |
| Peanut oil | Sigma-Aldrich | Cat# P2144 |
| Phosphate Buffer (0.2 M, pH 7.4) | Moltox | Cat# 26-510.047 |
| Polidocanol (nonaethylene glycol monododecyl ether) | Sigma-Aldrich | Cat# P9641 |
| Progesterone | Sigma-Aldrich | Cat# P8783 |
| Sesame oil | Sigma-Aldrich | Cat# S3547 |
| Sodium Deoxycholate | Sigma-Aldrich | Cat# 30970 |
| Sulfo-cyanine3 azide | Lumiprobe | Cat# 1330 |
| THEED (N,N,N',N'-Tetrakis(2-hydroxypropyl) ethylenediamine) | Sigma-Aldrich | Cat# 122262-1L |
| TritonX-100 | Sigma-Aldrich | Cat# X100 |
| Tween-20 | Sigma-Aldrich | Cat# P9416 |
| Urea | Sigma-Aldrich | Cat# U5128 |
| **Critical commercial assays** | | |
| Hemoccult Guaiac Fecal Occult Blood Test Systems | Beckman Coulter | Cat# 60151A |
| **Experimental models: Organisms/strains** | | |
| Mouse: Amhr2$^{Cre}$: B6.129(Cg)-Gt(ROSA)26Sortm4(ACTB-t dTomato,-EGFP)Luo/J | Ron Chandler (Jamin et al.[57]) | N/A |
| Mouse: B6: C57BL/6 | The Jackson Laboratory | RRID: IMSR_JAX:000664 |
| Mouse: CD-1, vasectomized | Charles River | RRID: IMSR_CRL:022 |
| Mouse: Cxcl15$^{iCre}$ | Andrew Kelleher (Kelleher et al.[23]) | N/A |
| Mouse: GsD: Gt(ROSA)26Sortm1(CAG-Chrm3*/GFP,c AMPRE-luc)Berd | Bruce Morgan (Akhmedov et al.[58]) | N/A |
| Mouse: mTmG: B6.129(Cg)-Gt(ROSA)26Sortm4(ACTB-tdTomato,-EGFP)Luo/J | The Jackson Laboratory (Muzumdar et al.[55]) | RRID: IMSR_JAX:007676 |
| Mouse: nTnG: B6N.129S6-Gt(ROSA)26Sortm1(CAG-tdTomato*,-EGFP*)Ees/J | The Jackson Laboratory (Prigge et al.[56]) | RRID: IMSR_JAX:023537 |
| **Software and algorithms** | | |
| Fiji/ImageJ version 1.54p | ImageJ | https://imagej.net/software/fiji/ |

| | | |
|---|---|---|
| Prism version 11.0.2 | GraphPad | https://www.graphpad.com/features |
| Zeiss Arivis Pro | Zeiss | https://www.zeiss.com/microscopy/us/products/software/arivis-pro.html |
| Leica Application Suite X (LAS X) software | Leica | https://www.leica-microsystems.com/products/microscope-software/p/leica-las-x-ls/ |
| NIS-Elements version 5.42.06 | Nikon Instruments | https://www.microscope.healthcare.nikon.com/products/software/nis-elements |
| **Other** | | |
| CFI60 Plan Apochromat Lambda 20×/0.75 NA objective lens | Nikon Instruments | Cat# MRD00205 |
| Compresstome® (vibrating microtome) | Precisionary Instruments | Cat# VF-510-0Z |
| Cryostat CM 1860 | Leica | N/A |
| CSU-W1 spinning disk confocal system | Yokogawa | Cat# 99927 |
| DM6 B upright microscope | Leica | https://www.leica-microsystems.com/products/light-microscopes/p/dm4-b-and-dm6-b-upright-microscopes/ |
| EC Plan-Neofluar 5x/0.16 NA nD=1.33-1.58 objective lens | Zeiss | N/A |
| Hydrophobic barrier pen | Vector Laboratories | Cat# H-4000 |
| Inverted Ti-2E microscope | Nikon Instruments | Cat# MEA54000 |
| K8 camera | Leica | https://www.leica-microsystems.com/products/microscope-cameras/p/k8/ |
| ORCA-Fusion Gen-III sCMOS camera | Hamamatsu | Cat# 77054115 |
| ProLong™ Diamond Antifade Mountant | Invitrogen | Cat# 36961 |
| ProLong™ Gold Antifade Mountant | Thermo Fisher Scientific | Cat# P36930 |
| Somnosuite (low-flow anesthesia system) | Kent Scientific | Cat# SS-01 |
| Superfrost Plus slides | Thermo Fisher Scientific | Cat# 22037246 |
| Tissue-Tek® OCT compound | Sakura | Cat# 4583 |
| VT1000 S Vibratome | Leica | RRID: SCR_016495 |
| Lightsheet.7 microscope | Zeiss | N/A |
